# Development of sound production in *Danionella cerebrum*

**DOI:** 10.1101/2024.03.25.586601

**Authors:** Antonia H. Groneberg, Lena E. Dressler, Mykola Kadobianskyi, Julie Müller, Benjamin Judkewitz

## Abstract

Acoustic signaling, integral to intraspecific communication and reproductive behavior, undergoes notable changes during an animal’s ontogenetic development. The onset and progression of this maturation in fish remains poorly understood. Here, we investigate the ontogeny of acoustic communication in the micro glassfish *Danionella cerebrum* (DC), one of the smallest known vertebrates and an emerging model organism in neuroscience. Adult DC males produce audible clicks that appear in sequences with a repetition rate of 60 or 120 Hz, caused by consecutive unilateral or alternating bilateral compressions, respectively. To investigate the maturation of this ability, we performed long-term sound recordings and morphological studies of the DC sound production apparatus throughout its ontogenetic development. We found that DC start producing clicks during the second month of their lives and continually increase their abundance and structured repetition over the course of the following one to two months. The sound production machinery, including specialized bone and cartilage structures, start to form in male DC after approximately four weeks and prior to full maturation of the reproductive organs. While the DC clicks increase in amplitude with age and body size, click repetition rates of 60 and 120 Hz are stable throughout development. This suggests a fully developed central pattern generator in juvenile males, yet a continued maturation of the drumming apparatus capable of creating louder sounds.

## Introduction

Acoustic communication can be found throughout the animal kingdom in rich spectral and temporal diversity. While the mechanisms of sound production are manifold, from bone stridulation to larynx vibrations (Suthers et al., 2016), a common theme is that sounds are associated with reproductive and territorial behavior.

In sound producing species with parental care, acoustic signals are typically produced already in early developmental stages, for instance as retrieval calls upon maternal separation (i.e. in bats (Bohn et al., 2007), squirrel monkeys (Symmes & Biben, 1985), rats (Brudzynski et al., 1999) and seals (Sauvé et al., 2015) or as food calls of nestlings (Briskie et al., 1999). Vocalizations change throughout the lifespan of an animal, especially around the time of sexual maturation. A well-known example is the change in pitch in human males during puberty. In many vocal learner species, individuals acquire a new vocal repertoire upon sexual maturation, as prominently studied in songbirds (Nottebohm, 1970). The trajectory of sound development of vocal learners can vary. For instance, in the greater sac-winged bat, pups already start vocal imitation of male territorial song (Knörnschild et al., 2010), while pups of neotropical singing mice are highly vocal right after birth, but stop vocalizing before weaning and then produce their mature advertisement song *de novo* (Campbell et al., 2014).

In soniferous egg-laying species without parental care, acoustic communication is present in adults and tends to commence in juvenile stages. For instance, tadpoles do not emit sounds, but vocalizations are found in juvenile spadefoot toads (Ten Hagen et al., 2016). In several species of sound producing Sciaenidae fish, the sound generating muscles only form in juvenile fish in parallel with gonad development (Hill et al., 1987). In contrast, it has been reported that coral reef gray snapper already produce nocturnal sounds as pre-settlement larvae with a body length of a mere centimeter (Staaterman et al., 2014). This suggests that within the teleostean fishes, there is variability when and how the sound production systems develop in an animal’s lifetime. In addition, there is limited information to date linking the morphological ontogenesis of the sound producing organs to sound production and resulting sound characteristics across development.

Here, we investigate the interplay between morphological ontogenesis and the development of sound production in the transparent miniature teleost *Danionella cerebrum* (DC). DC has an adult body length of only ∼12 mm and the smallest known vertebrate brain (Britz et al., 2021; Penalva et al., 2018; Schulze et al., 2018). Its small size, together with its life-long transparency has made DC an emerging model system for neuroscience, suitable for whole-brain optical measures of neuronal activity (Schulze et al., 2018; Kadobianskyi et al., 2019; Hoffmann et al., 2023; Lam, 2022; Zada et al., 2023; Lee & Briggman, 2023). Adult DC have a male-specific trait of acoustic signaling, producing click-like sounds by drumming a piece of cartilage against the anterior swim bladder (Schulze et al., 2018; Cook et al., 2024; Vasconcelos et al., 2024). The highly specialized drumming apparatus is only present in male DC and absent in females. To date it is unknown when DC develops its sexually dimorphic sound production apparatus and at which developmental stage acoustic signaling begins. We addressed these questions through a combination of longitudinal sound recordings and morphological comparisons.

## Results

### First clicks appear in juvenile *Danionella cerebrum*

To probe at what developmental stage DC first emit sounds, we raised a group of 100 fish in a stand-alone 30×60 cm aquarium and performed underwater sound recordings once a week. The first time point at which click sounds were clearly distinguishable from background noise was 6 weeks post fertilization. From 6 to 8 weeks of age we observed an increase in the abundance, as well as the amplitude of clicks (example sound traces shown in Fig. 1A).

**Figure 1:**
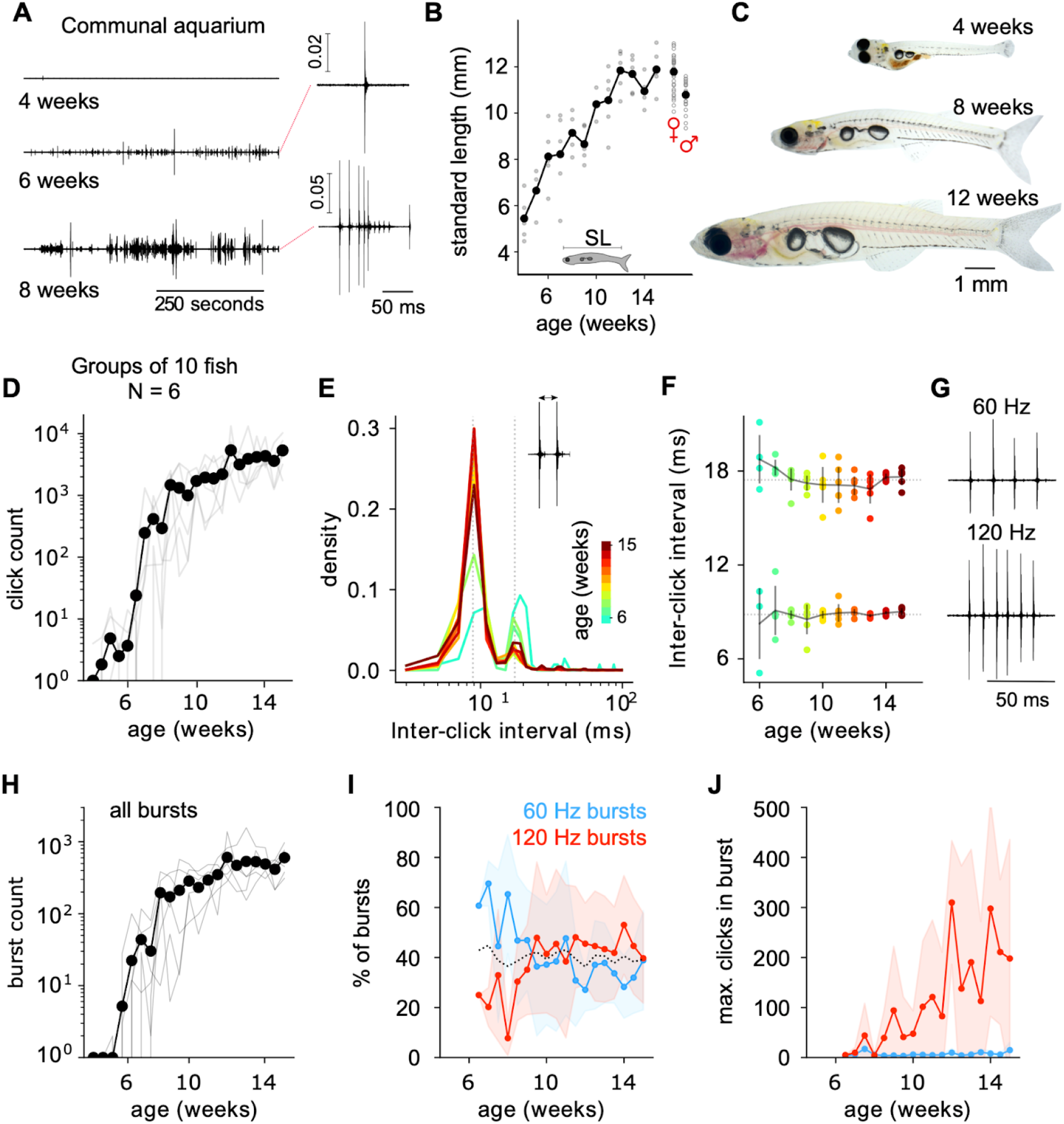
Emergence of click sounds in juvenile DC. **A**) Example sound traces from hydrophone recordings across different stages of development in a communal aquarium containing 100 fish. **B**) Standard length measures of DC over the course of three months, as five samples per age (shown as gray dots) and mean between samples (connected black dots). At 15 weeks-post fertilization female DC are larger than males. **C**) Example images of larval, juvenile and adult DC according to stated age and scale bar. **D-J** Six groups of 10 DC were raised in individual tanks and recorded twice a week with a hydrophone. **D**) Number of clicks detected per recording time point, shown as connected lines for each recording group (gray lines) and mean across groups (black dots and line). **E**) Inter-click interval refers to the time between two adjacent clicks (schematic in inset). Density distribution of inter-click intervals color-coded for ages 6 to 15 weeks. **F**) The peaks of the bimodal distribution in E are shown for each recorded age. Circles show each recording group, color-coded by age as in E, and black lines and error indicate mean ± SD values across recording groups per age. Dashed, horizontal lines indicate means across age and groups (8.8 ± 0.8 and 17.4 ± 1.0 ms). **G**) Example sequences of clicks with inter-click intervals of 60 and 120 Hz. **H**) Number of bursts detected per recording time point. Bursts are defined as groups of consecutive clicks with a max. interval of 25 ms. Mean across recording groups and individual recording groups are shown as black dots and gray lines, respectively. **I**) The percentage of 60 and 120 Hz bursts of all detected bursts per recording point, shown as blue and red mean values with shaded SD, respectively. In total these single frequency burst types make up approximately 80 % of all bursts across age (dashed black line). **J**) The maximum number of clicks detected in a single burst shown across age for 60 and 120 Hz bursts as blue and red mean values with shaded SD.

To examine sound production in more details across age, we next raised six groups of ten fish in 10x10 cm tanks, equipped with one hydrophone each, and recorded twice a week over the course of 11 weeks starting at 4 weeks after fertilization. To accurately measure the growth rate without disturbing the sound recording tanks, we raised additional individuals under identical conditions and measured the standard length (SL, distance from snout to tail-base). DC grew over the course of three months until they reached a SL of 11.9 ± 1 mm (Fig. 1B, N=5 per age). These measurements were drawn from a random pool of a mixed sex fish population. Females are generally larger than males at 15 weeks post fertilization (11.8 ± 0.7 and 10.8 ± 0.6 mm, respectively, two-sided t-test: t=5.3, p<0.001, N=37 females, N=23 males). Growth in body size was accompanied by gross morphological changes, including a change in the shape of the swim bladder complex (Fig. 1C).

Similar to the communal aquarium recording, the number of recorded clicks increased between 6 and 9 weeks post fertilization (Fig. 1D), corresponding to 8.1 ± 1.3 mm SL at 6 weeks and 9.1 ± 0.8 mm SL at 9 weeks (Fig. 1B). The exact age when groups started producing clicks varied between 6.5 and 8 weeks in the different groups (see gray lines in Fig. 1D). The average age with more than 20 recorded clicks per group was 7 ± 0.5 weeks.

### Bursts of clicks appear at stable repetition rates and become longer with age

Adult DC clicks appear with a repetition rate of either 60 or 120 Hz (Schulze et al., 2018; Vasconcelos et al., 2024), corresponding to repeated unilateral and alternating bilateral contractions of the drumming muscles (Cook et al., 2024). Both click repetition rates were present in juvenile DC from the earliest sound-producing age on (Fig. 1E) and the peak of the bimodal distribution did not change with age (Fig. 1F). Groups of repeated clicks, hereafter referred to as bursts, tend to occur with a single click repetition rate of either 60 or 120 Hz (example bursts shown in Fig. 1G). Similar to the total number of clicks (Fig. 1D), the number of bursts increased between 6 and 9 weeks post fertilization (Fig. 1H). In total, single-frequency bursts make up approximately 80% of all detected bursts throughout all ages tested. At younger ages, DC tend to produce more 60 Hz bursts, while at later stages both frequencies occur at a similar rate (Fig. 1I). Finally, we measured the length of each burst as the number of clicks in the sequence (Fig. 1J). The maximum number of clicks observed in a burst reached up to several hundred for 120 Hz bursts in older fish, while for 60 Hz bursts it did not surpass ten.

### Clicks become louder with age while preserving the sound profile

In the communal aquarium recordings (Fig. 1A), as well as the group recordings (Fig. 2A), the amplitude of the clicks increased with age. Because clicks are generated by compressing the anterior swim bladder (Cook et al., 2024) and the swim bladder size increases with age (Fig. 2B), we next asked how click features change as a function of swim bladder size. Since in the previous sound recordings the distance of the sound source to the hydrophone was unknown, the sound attenuation and thus the actual sound volume could not be reliably established. To obtain recordings of clicks at a controlled distance, we set up groups of three age-matched DC juvenile or adult males in a small compartment surrounded by five hydrophones at a distance of 3.5 cm to the center. In this single measure design the fish were raised in the fish facility under circulating water and had a faster growth rate. In order to compare the developmental stage to the previous experiments we color-coded the age according to the body length and size of the anterior swim bladder. Note that the oldest juveniles tested in this setup had the same SL as the adult group (10.5 mm), but that adult DC had a larger swim bladder diameter along the dorsoventral axis (hereafter referred to as height, Fig. 2B inset on the right). The distance-controlled click recordings allowed us to compare the click profile, which was stereotyped within each developmental group (Fig. 2C). While the shape of the pulse was similar across age, with its peaks aligned in time, peak amplitude increased with age (Fig. 2C, right). Indeed, click amplitude was positively correlated with swim bladder height (Fig. 2D; Pearson r=0.99, p=0.013, N=4; one data point per age), and reached up to 140 dB, matching previous reports of DC sound pressure levels (Cook et al., 2024). In contrast, the peak-to-peak time of the main click pulse was not correlated with swim bladder height (Fig. 2E; Pearson r=0.20, p=0.80). This suggests that the sounds of individual clicks are stable, but get louder with age.

**Figure 2:**
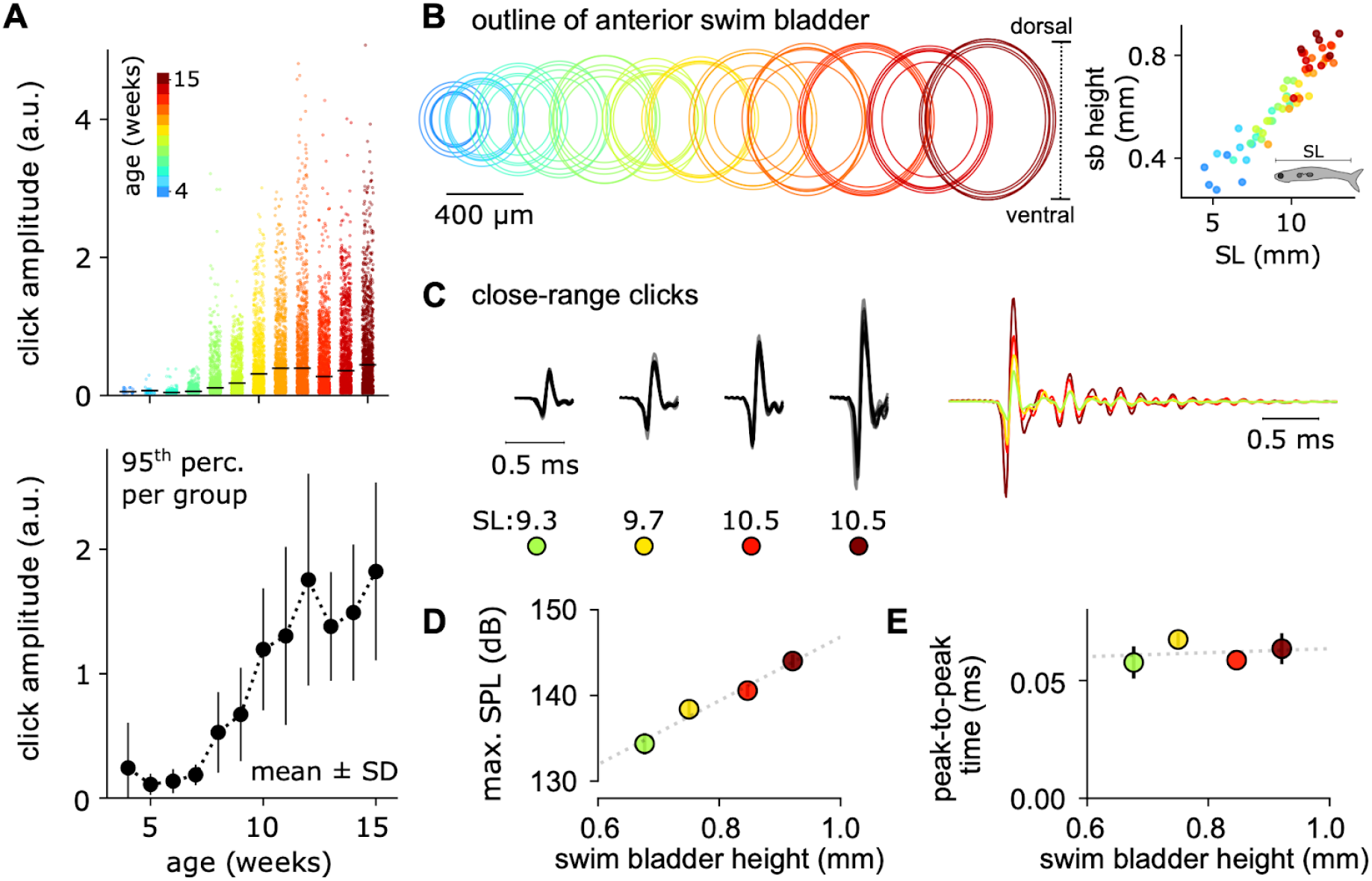
DC sound features across age. **A**) Click amplitude in arbitrary units (a. u.) across age, measured in six groups of 10 DC as a repeated measure. Top: 200 randomly selected clicks from all recording groups, color-coded by age, with median values as black lines. Bottom: the loudest 95th percentile of all recorded clicks per group is shown as the mean and standard deviation between the six recording groups. **B**) Outline of the anterior swim bladder according to the shown scale, color-coded for age as in A. Inset on the right shows the standard length against swim bladder (sb) height, measured as the dorsal to ventral diameter, for each measure per age. Note that this data was collected from fish that were raised in parallel with the sound-recorded groups under the same conditions and sampled at a weekly rate as single measures, not repeated measures. **C-E** Groups of three males at different developmental stages we recorded in a small enclosure surrounded by five hydrophones at a distance of 3.5 cm. Because growth rates depend on the raising conditions, the relative age is color-coded according to the measured standard length (SL in mm) and dark-red for adult males. **C**) Sound amplitude profiles of the ten loudest clicks recorded per developmental time point. Profiles of a single click per developmental stage are overlaid in color on the right. Note that red and dark red samples have the same standard lengths, but stem from juvenile and adult samples, respectively. **D**,**E** Maximum click amplitude converted to sound pressure level (**D**) and peak to peak time of the main sound pulse (**E**) is plotted against the swim bladder height of the recorded DC males. Shown are the mean values ± standard deviation of the ten clicks shown in C. Dashed lines show the slope of a Pearson correlation. Mean values for each recorded group were used for the correlation to avoid pseudoreplication. Pearson r = 0.99, p = 0.01 for SPL and r = 0.20, p = 0.80 for peak-to-peak times.

### Drumming apparatus begins to form and matures in juvenile males

We next investigated the development of the DC sound production apparatus via bone and cartilage staining. In adults the drumming apparatus and the skeletal structures associated with the anterior swim bladder are sexually dimorphic (Fig. 3 A,B). Adult males have a hypertrophied 5^th^ rib, a globular cartilage and two extra components of the os suspensorium (OS), a bone structure that supports the anterior swim bladder anteriorly and dorsally (Britz et al., 2021). The inner part of the male OS (iOS) has a posterior extension on the dorsal side that extends over the swim bladder. The male outer OS (oOS) extends further rostral and has an additional connection to the transverse process of the second vertebra. An additional feature of the DC sexual dimorphism is the position of the vent of the digestive tract, which is shifted anteriorly to the pelvic fin in males (Britz et al., 2021). This rerouting of the vent is the last sexually dimorphic developmental change in juvenile males, occuring after all components of the drumming apparatus have developed (Fig. 3A,B, bottom). However, the connecting flanges between the outer and inner arms of the OS, as well as the connection between the oOS and the spine continue to grow in adult males.

**Figure 3:**
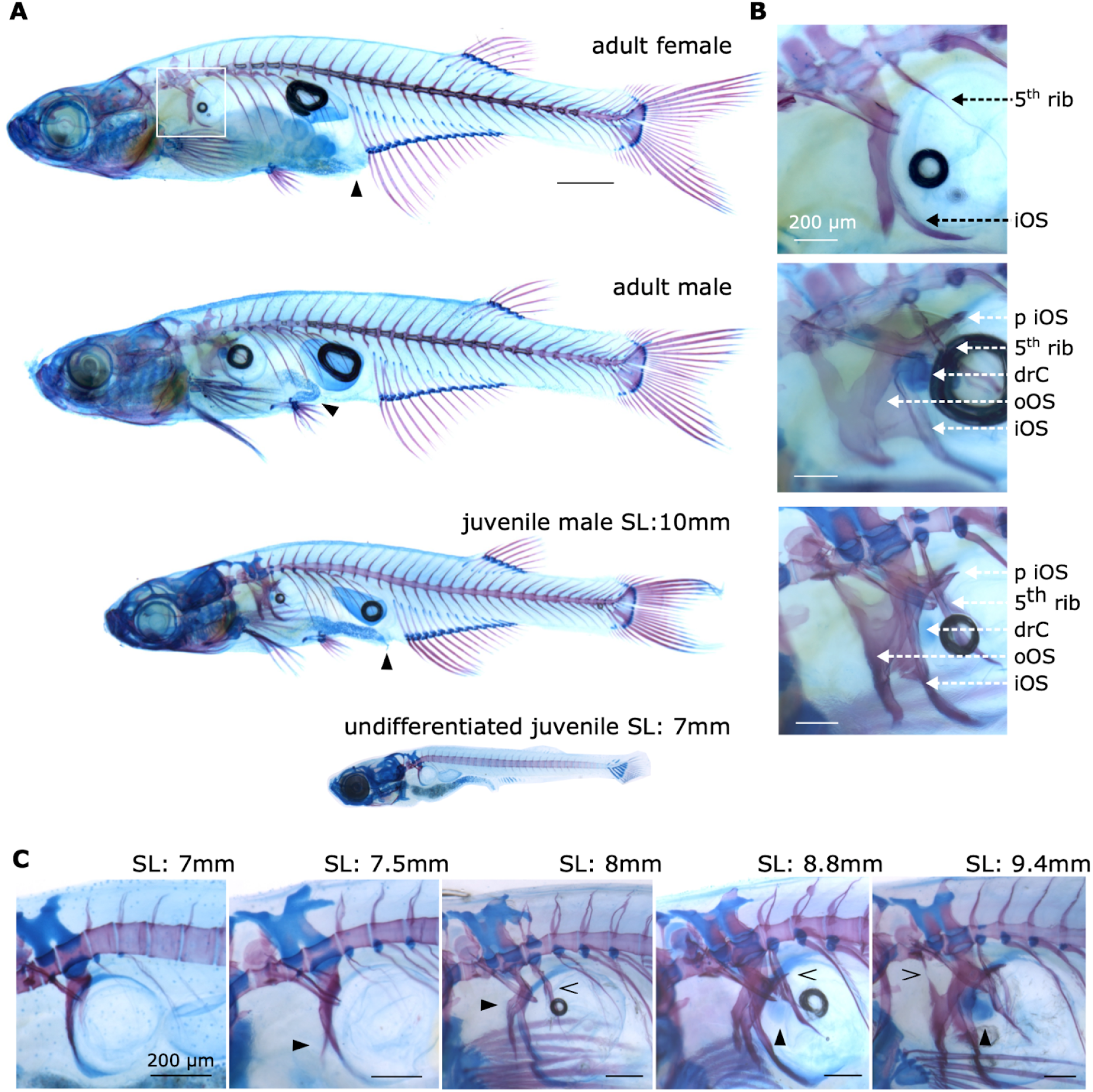
Development of the sound production apparatus in DC males. Bone and cartilage staining via Alizarin Red S and Alcian Blue, respectively, in unpigmented DC (tyrosinase knock-out). **A**) Exemplary adult female, adult male, juvenile male and an undifferentiated juvenile DC. Black arrowheads point to the anal exit of the gastro-internal tract, which is rerouted in adult males, but not in juvenile males and females. Scale bar 1 mm. **B**) Zoom-in on the anterior swim bladder region in the samples shown in A (white box on female shows zoom-in region). **C**) Zoom-in on the anterior swim bladder region in DC juveniles atdifferent developmental stages. Standard length (SL) as stated in the figure. Below a SL of 8 mm, the sex could not be determined. SL 8mm and above are clearly identified male samples. Black arrowheads point towards: the developing oOS in SL 7.5 mm and 8 mm and the developing drumming cartilage in SL 8.8 mm and. 9.4 mm. Open arrowheads point towards the 5^th^ rib in SL 8 mm and 8.8 mm; and to the connection of the oOS to the spin in SL 9.4 mm. Abbreviations: inner arm of the os suspensorium (iOS), posterior extension of the iOS (p iOS), outer arm of the os suspensorium (oOS), drumming cartilage (drC). The black circles seen in some of the images are the remainder of the partially deflated swim bladder. SL was measured post-mortem, prior to fixation, to avoid distorted measures due to shrinkage.

In 2–4-week-old DC, sexes cannot yet be distinguished morphologically. Undifferentiated juveniles resemble the adult female morphotype in their skeletal anatomy, with a thin 5^th^ rib and only the lower extension of the iOS present (Fig. 3C, SL:7mm). At 7.5mm the still undifferentiated juvenile develops the short, female version of the oOS (Fig. 3C, SL:7.5mm). The first male-specific component that emerges at a SL of 8.0 mm is a hook-like outgrowth of the oOS, which starts to form around the same time of the ossification of the 5^th^ rib (Fig. 3C, SL:8mm). The next component is the posterior extension of the iOS over the swim bladder, which coincides with a faint first staining of the drumming cartilage (Fig. 3C, SL:8.8mm). The final step in the emergence of the male drumming apparatus is the connection of the oOS back to the spine (Fig. 3C, SL: 9.4mm). The oOS continues to grow and becomes increasingly ossified during further maturation (Fig. 3A,B). In summary, the drumming apparatus components form in juvenile males at a SL between 8 and 10mm. This is inline with the ontogeny of recorded clicks sounds in our behavioral experiments (Fig 1 and 2).

## Discussion

*Danionella cerebrum* is a miniature fish with a male-specific trait of acoustic signaling. Here we showed that DC males start producing sounds after 1 to 1.5 months of development, at a SL of ca. 9 mm. In line with this, we found that the sound-producing drumming apparatus develops in male specimens between 7 and 10 mm of SL. A full set of drumming components is present at approximately 9 mm SL, while the OS continues to ossify and the rerouting of the vent is yet to occur. We also found that click repetition rate and broadband frequency spectra were consistent across age, while sound intensity increased with maturation.

### Changes in click characteristics across development

Acoustic allometry, a relationship between body size and sound features, has been described across species of primates and carnivore mammals (Bowling et al., 2017), as well as anurans (Gingras et al., 2013). While within-species allometry is more complex in vocal learning species and possibly species with the capability of volitional sound modulation (Garcia & Ravignani, 2020), it has been described in several fish species with differing sound production mechanisms. For instance, larger fish of the cichlid species *Metriaclima zebra* produce lower frequency and louder sounds (Bertucci, Attia, et al., 2012), a trend that was also found when comparing juvenile and adult fish (Bertucci, Scaion, et al., 2012). Similar results have been reported for Lusitanian toadfish *Halobatrachus didactylus* (Vasconcelos & Ladich, 2008), grey gurnard *Eutrigla gurnardus* (Amorim & Hawkins, 2005), mochokid catfish *Synodontis schoutedeni* (Lechner et al., 2010) and croaking gourami *Trichopsis vittata* (Wysocki & Ladich, 2001). A study performed on the weakfish *Cynoscion regalis* described a negative correlation between pulse duration and dominant frequency (Connaughton et al., 2000). By comparing individual sizes, the authors found that larger males produce louder sounds with a longer pulse duration and lower dominant frequency, without any clear change in pulse repetition rate. This is mostly in line with our results in DC, where we found an increase in sound amplitude with size and age (Fig. 2A,D), yet no change in pulse repetition rate (Fig. 1F). Unlike in weakfish and other species, we found no evidence of changes in pulse duration or dominant frequency with size and age (Fig. 2C,E). Most sonic fish with a swim bladder mechanism, including the weakfish, use direct muscle contractions to produce sounds. In contrast, DC employ an indirect sound production mechanism where a drumming cartilage snaps out and strikes the swim bladder upon muscle contraction (Cook et al., 2024). Therefore, unlike in weakfish, the strength or size of the sonic muscle in DC seems less likely to influence pulse duration and thus dominant frequency. Here we show a positive correlation between sound amplitude and swim bladder height (Fig. 2D), while the temporal click profile remained similar (Fig. 2C,E). The correlation between swim bladder height and click amplitude is unlikely to be a causal link, but rather a byproduct of increased maturity of the structures of the drumming apparatus, such as the inner arm of the OS with its dorsal and ventral extension around the swim bladder and likely the mass of the sonic muscle. In line with this, a study comparing sonic muscle mass in the weakfish across spawning seasons showed a positive correlation with sound pressure levels (Connaughton et al., 1997).

### Maturation of sound structure in DC

Adult DC males produce clicks at repetition rates of 60 and 120 Hz by repeated unilateral or by alternating bilateral contractions, respectively (Cook et al., 2024). The presence of both repetition rates in juvenile DC suggests that the pattern generating neuronal circuitry that controls the alternating contractions is fully functional at the developmental onset of acoustic signaling. Similar results have been reported in weakfish, where pulse repetition rate did not differ with body size (Connaughton et al., 2000). Our dataset shows a trend for older DC to have proportionally more clicks at 120 Hz, as seen in the higher peaks of the histogram (Fig. 1E). This can be explained by the increasing length of 120 Hz bursts with age of up to multiple hundred clicks. 60 Hz bursts in contrast do not get longer than 10 clicks (Fig. 1J). Furthermore, we show that while a stable 80% of all bursts are single-frequency bursts throughout all ages tested, younger ages show a higher likelihood of 60 Hz bursts over 120 Hz (Fig. 1I). We note that the high variability in burst types and durations between groups could be a result of not having the same sex ratio in each tank, due to the repeated-measure design of the experiment. Given the positive relationship between maturation and sound amplitude, there is a possibility that younger fish produced sounds below the detection threshold of our setup. However, since our morphological analysis showed that the first cartilage staining appeared in fish with a SL of at least 8 mm (matching an age of 6-7 weeks in the repeated measure sound recordings), it is unlikely that DC is able to produce sounds earlier than that. Indeed, in the repeated measure experiments first clicks appeared at 8.1 ± 1.3 mm SL (Fig. 1B and D). Since there is individual variability in growth rate, the larger males in the group recordings could have been the ones first producing sounds. In the close-range click recordings with fewer fish per group and only males, body size measurements align well with the morphological results; the youngest clicks we recorded were at 9.3 mm SL (Fig. 2C), a stage at which all components of the drumming apparatus have formed (Fig. 3C).

### Development of sound-related anatomy

To date there is limited knowledge about the link between morphology of the sound producing machinery and resulting sound features across ontogenic development. Several studies have measured sound production throughout development in fish (Vasconcelos & Ladich, 2008, Amorim & Hawkins, 2005, Lechner et al., 2010), yet without morphological measures beyond body size. In contrast, few studies have described the morphological development of sound-related structures (Hill et al., 1987, Fine, 1989, Brantley et al., 1993), however, without measuring sound output. One exception is a study by Kéver and colleagues in *Ophidion rochei*. Unlike female DC, females of *O. rochei* have a less developed version of the male sound apparatus and can produce sounds. The first juvenile sounds that were recorded in *O. rochei* resembled female sounds rather than adult male sounds (Kéver et al., 2012). Similar to DC, the sound apparatus of *O. rochei* is sexually dimorphic and juvenile males initially resemble the female morphotype and then differentiate new structures during gonadal maturation (Kéver et al., 2012). While we did not measure gonadal maturation, we noticed that the rerouting of the gut vent appeared to be the last morphological change in male DC. During the close-range click experiments, we noticed that we were only able to obtain clicks from groups of juveniles in which at least one of the males had a rerouted vent. Although merely anecdotal at this stage, this observation hints at a sound production onset after full sexual differentiation.

### *Danionella* as a model for development of acoustic communication

To date our understanding of the neuronal processes underlying the ontogenic development of sound production and acoustic communication is limited due to methodological constraints of large-scale neuronal recordings across different developmental stages. DC, with optical accessibility during its entire life, offers a unique opportunity as a vertebrate model to study the ontogeny of acoustic communication. An essential part of studying the development of acoustic communication is sound perception. Several studies have applied auditory brainstem recordings to probe hearing capabilities across age and related it back to the sound parameters individuals were able to produce (Lechner et al., 2010; Vasconcelos & Ladich, 2008; Wysocki & Ladich, 2001). The small, transparent DC will enable whole-brain imaging of sound perception across development to establish not only when sounds are transduced to neural signals in the brainstem, but also when they start to trigger activity in brain regions that are associated with social behaviors across vertebrates, such as the preoptic area of the hypothalamus (Goodson, 2005). Notably, the Weberian apparatus shows an accelerated development in the Danionella species (Conway et al., 2021) and our morphological results also showed its fully ossified components, i.e. tripus and scaphium, prior to the development of any of the sexually dimorphic components of the sonic apparatus (Fig. 2C). This suggests that hearing develops prior to sound production in DC and it will be an interesting future study to measure the perception of social sounds in non-sonic juveniles in the future. Furthermore, an ancestrally shared development of the neuronal vocal-motor circuits between fish and vocal terrestrial mammals has been proposed (Bass et al., 2008). DC may therefore serve as a suitable model to study both the development of sound production circuitry, as well as auditory perception of acoustic social signals. Our study sets the stage for such investigations with the report of how and when DC first start producing sounds.

## Methods

### Animal husbandry

*Danionella cerebrum* were housed in water circulated Tecniplast aquaria with artificial fish water (Instant Ocean at a conductivity of 350 µS/cm, adjusted to a pH of 7.3 using NaHCO_3_) at 27 °C and a 14:10h light-dark cycle. For all behavioral experiments wild type (WT) DC were used. For morphological stainings a tyrosinase knock-out line (*tyr-/-*) was used to remove melanin pigmentation for increased transparency to see bone and cartilage structures.

### Raising conditions

Fertilized eggs from communal breeding in the regular home tanks were collected and kept in 10 cm petri dishes containing embryo water (600 mg/L Instant Ocean, 150 mg/L NaHCO_3_, 50 mg/L CaCl_2_ 2H_2_O, 0.5 mg/L Methylene Blue) at a density of < 20 eggs per dish. At 5 days post fertilization (dpf) larvae were transferred to 1 L plastic tanks (Tecniplast breeding tank) with A2000 water (2 g/L Instant Ocean - Aquarium system, 150 mg/L NaHCO_3_, pH 7.4, conductivity 1.5 mS/cm) and L-type rotifers (Brachionus plicatilis, planktovie.biz). For regular facility rearing, larvae were transferred to the water circulation system at 10 dpf and fed twice daily with artemia, as well as dry food (Gemma micro 75). For rearing in experimental tanks, larvae were transferred at 3 weeks post fertilization (wpf) and fed once daily with artemia. Water was either circulated by a pump or manually exchanged twice a week in the small tanks.

### Repeated measure sound recordings

For the single recording of a large group (Fig. 1D) 100 larvae at 3 wpf were transferred to a standalone aquarium (30x60 cm) filled with 18 L artificial fish water with a circulating pump. Three times per week 1 L of tank water was replaced with fresh artificial fish water. Water temperature was maintained at 27 °C with three aquarium heater thermostats (Sera). Once a week in the morning between 9 and 11 am a hydrophone (H3x, Aquarian Audio) was inserted into the tank and 1 hour sound recordings were acquired with a datalogger (Zoom field recorder - Fn8) at a sampling rate of 48 kHz, while the circulating pump was switched off.

For the multiple group recordings (Fig. 1D-E) 10 larvae at 3 wpf were kept inside 10x10 cm tanks filled with 800 mL of artificial fish water (at 27 °C). This led to a higher fish density than in the large aquarium or the fish facility, but it was a tradeoff between recording quality and fish number. Because a higher fish density can lead to developmental delays, we measured body size in addition to biological age. For length measurements, several extra tanks were raised under the same conditions as the sound-recorded fish. 5 of these sample fish were collected per week, euthanised and photographed under a brightfield microscope (Olympus MVX-10 with a MV PLAPO 1X objective, connected to a FLIR usb color camera). Standard length measurements were manually performed post hoc on the collected images, as measured from snout to the base of the tail, excluding the tail fins.

To avoid disturbance of the fish prior to sound recordings, a hydrophone (AS-1, Aquarian Audio) was placed inside each tank the evening prior to a 2-hour recording from 9 to 11 am the following morning. This procedure was repeated twice a week over the course of 11 weeks, to record ages 4 through 15 weeks post fertilization. Sound recordings were obtained with a sampling rate of 51.2 kHz via a National Instruments data acquisition card (NI-9231) controlled through a custom written python script. Each AS-1 hydrophone was amplified with a PA-4 hydrophone preamplifier at an inverted wiring state (Aquarian Audio), powered by a 12V battery.

### Close-range click recordings

Three age-matched fish were placed in a 5x5x5 cm box made out of transparent acrylic and walls of blue cling film. This box was placed in a holder in the center of a 27x27x30 cm glass aquarium filled to ca. 27 cm. Five AS-1 hydrophones were positioned around the four vertical walls and the bottom of the inner box, with a fixed distance of 3.5 cm to the center of the box. Sound recordings were acquired for several hours at a sampling rate of 102.4 kHz using a National Instruments data acquisition card (NI-9250). Clicks that were detected on at least four hydrophones at coinciding or adjacent samples were considered centered clicks, produced by fish at the center of the box and hence an equal distance of ca. 3.5 cm. To control for the different amount of recorded center clicks per group, 200 center clicks were randomly selected. Out of those 200 clicks the single loudest was determined and its sound trace from the channel with maximum amplitude used for further analysis. The fish used in this experiment were raised in the facility and had a faster growth rate than the repeated-measure experimental fish. Therefore, the fish age is expressed in standard length (SL) as measured after the sound recordings and expressed as the mean between the three fish of each group.

The matching between biological ages, SL and swim bladder height (Sb-height) were: 5 week old fish SL = 9.3 ± 0.4 mm, Sb-height = 0.68 ± 0.04 mm; 6 week old fish SL = 9.7 ± 0.6 mm, Sb-height = 0.75 ± 0.06 mm; 8 week old fish SL = 10.5 ± 0.2 mm, Sb-height = 0.85 ± 0.03 mm and adult (older than 3 months) SL = 10.5 ± 0.6 mm, Sb-height = 0.92 ± 0.07 mm.

### Click detection

Clicks were detected with a custom-written python code. Waveform traces were bandpass-filtered between 3 and 20 kHz and peaks were detected using the waveform-envelope and a refractory period of 5 ms between peak times. Filtering parameters and detection thresholds were always kept constant for all recorded ages within one experiment. Bursts were defined as sequences of clicks appearing at less than 25 ms between two clicks.

### Morphological measures

*tyr-/-* specimens of different ages were euthanized in ice water and fixed overnight in 4% paraformaldehyde in 1x PBS at 4 °C. After brief washing in 1x PBS, samples were stained for 24 hours with a double staining solution of 0.001% Alcian Blue (Roth) and 0.005% Alizarin Red S (Sigma-Aldrich) with 200mM MgCl_2_ x6 H_2_O in 70% EtOH (Walker & Kimmel, 2007). After two brief washes in pure H_2_O, samples were kept in 20% glycerol with 0.25% KOH for 2 hours and 50% glycerol with 0.25% KOH overnight. Thereafter, samples were kept in a commercial refractive index matching solution (RI=1.465, EasyIndex Life Canvas) for 2-3 days and imaged under a brightfield illuminated microscope (Olympus MVX-10). All steps after fixation were performed at room temperature. Standard length measurements were taken post-mortem prior to fixation to account for possible shrinkage due to sample dehydration.

## Acknowledgements

We thank Johannes Veith, Maximilian Hoffmann, Jörg Henninger and Verity Cook for assistance with methodological implementation and discussions of the ongoing work. We also thank our fish facility team for excellent fish care and experimental support. We thank Ralf Britz and Andrew Bass for feedback on the manuscript. This work was supported by the German Research Foundation (DFG, projects EXC-2049-390688087 and 432195732) the European Research Council (ERC2021-CoG-101043615), the Einstein Foundation (EPP-2017-413) and the Alfried Krupp Foundation. A.G. acknowledges funding by European Union’s Horizon 2020 research and innovation programme under the Marie Skłodowska-Curie actions, grant agreement number H2020-MSCA-IF 101026592.

## Author contributions

Conceptualization: AG, BJ

Methodology: AG, LD, MK, JM

Investigation: AG, LD, MK, JM

Visualization: AG

Funding acquisition: BJ, AG

Supervision: BJ

Writing – original draft: AG, BJ

Writing – review & editing: AG, LD, MK, JM, BJ

